# IL-15 complex-induced IL-10 enhances *Plasmodium*-specific CD4^+^ Tfh differentiation and antibody production

**DOI:** 10.1101/2023.10.06.561264

**Authors:** Morgan Bravo, Thamotharampillai Dileepan, Molly Dolan, Jacob Hildebrand, Jordan Wolford, Sara E. Hamilton, Anne E. Frosch, Kristina S. Burrack

## Abstract

Malaria, which results from infection with *Plasmodium* parasites, remains a major public health problem. While humans do not develop long-lived, sterilizing immunity, protection against symptomatic disease develops after repeated exposure to *Plasmodium* parasites and correlates with the acquisition of humoral immunity. Despite the established role antibodies play in protection from malaria disease, dysregulated inflammation is thought to contribute to the sub-optimal immune response to *Plasmodium* infection. *Plasmodium berghei* ANKA (PbA) infection results in a fatal severe malaria disease in mice. We previously demonstrated that treatment of mice with IL-15 complex (IL-15C; IL-15 bound to an IL-15Rα-Fc fusion protein) induces IL-10 expression in NK cells, which protects mice from PbA-induced death. Using a novel MHC class II tetramer to identify PbA-specific CD4^+^ T cells, herein we demonstrate that IL-15C treatment enhances Tfh differentiation. Moreover, genetic deletion of NK cell-derived IL-10 or IL-10R expression on T cells prevents IL-15C-induced Tfh differentiation. Additionally, IL-15C treatment results in increased anti-PbA IgG antibody levels and improves survival following reinfection. Overall, these data demonstrate that IL-15C treatment, via its induction of IL-10 from NK cells, modulates the dysregulated inflammation during *Plasmodium* infection to promote Tfh differentiation and antibody generation, correlating with improved survival from reinfection. These findings will facilitate improved control of malaria infection and protection from disease by informing therapeutic strategies and vaccine design.

## INTRODUCTION

*Plasmodium* parasites cause malaria in nearly 250 million persons per year, leading to >600,000 deaths annually worldwide. Protection from clinical disease develops following repeated infections and is mediated through humoral immunity. However, humans do not develop long-lived, sterilizing immunity (1–4). Indeed, antibody responses to *Plasmodium* antigens are short-lived and rapidly lost in the absence of continued parasite exposure (5–10). Dysregulated inflammation is thought to contribute to the short-lived response to *Plasmodium* (11–16). The *Plasmodium berghei* ANKA (PbA) mouse model recapitulates the dysregulated inflammation and resulting sub-optimal immunity seen in human malaria (12, 17).

Achieving high levels of antibody depends on antigen-specific B cells that, upon antigen encounter, proliferate and/or terminally differentiate into plasma cells or memory B cells, which seed the bone marrow and provide a lasting source of serum antibody. Effective induction of humoral immunity to most pathogens and vaccines requires help from CD4^+^ T helper (Th) cells, which are activated by pathogen-specific peptides presented on MHC class II molecules. CD4^+^ T cells can differentiate into several functionally distinct subsets, including type 1 (Th1), identified by expression of the transcription factor T-bet, and follicular helper (Tfh) cells, identified by expression of CXCR5, PD-1 and Bcl-6. In germinal centers (GCs), Tfh cells promote antibody responses, especially the generation of long-lived plasma cells and memory B cells during infections (18), including malaria (19, 20).

The study of *Plasmodium*-specific CD4^+^ T cell responses has been hampered by the lack of defined parasite-derived T cell epitopes. To investigate antigen-specific CD4^+^ T cells in the context of *Plasmodium* infection, groups have previously utilized surrogate activation markers (14), transgenic parasites expressing model antigens (21, 22), or TCR transgenic mice specific for *Plasmodium* antigens (23, 24). We utilized newly identified immunodominant CD4^+^ T cell epitopes (25) and generated a pMHCII tetramer to identify endogenous *Plasmodium*-specific CD4^+^ T cells in mice. Using a sensitive pMHCII tetramer-based cell enrichment method (26), we can identify CD4^+^ T cells specific for a peptide derived from ETRAMP (early transcribed membrane protein) within *Plasmodium* and presented in the context of I-A^b^ in C57BL/6 mice. This new reagent has enabled us to define the phenotype of *Plasmodium*-specific CD4^+^ T cells in mice and test strategies to modulate this response.

We previously showed that IL-15 complex (IL-15C: IL-15 combined with IL-15Rα) treatment prevents PbA-induced experimental cerebral malaria disease symptoms and death (27). IL-15 stimulates activation and expansion of natural killer (NK) cells and CD8^+^ T cells. Interestingly, we found that IL-15C treatment induces IL-10 expression from NK cells, which dampens the pathologic CD8^+^ T cell response (27). Using our novel CD4^+^ T cell tetramer to identify *Plasmodium*-specific CD4^+^ T cells in mice, we defined the phenotype of *Plasmodium*-specific CD4^+^ T cells and investigated the impact of NK cell-derived IL-10 on CD4^+^ T cell differentiation. Using conditional genetic knockout mice, we identified a novel pathway whereby NK cell-derived IL-10 acts directly on CD4^+^ T cells to promote Tfh differentiation, resulting in increased antibody formation during PbA infection and improving survival following reinfection.

## MATERIALS AND METHODS

### Mice

All strains are on the C57BL/6 background. C57BL/6 mice (strain #000664) were purchased from Jackson Laboratories and bred in-house. CD4-Cre mice (strain #022071) and *Il10ra*^fl/fl^ mice (strain #028146) were purchased from Jackson Laboratories. NKp46-iCre mice were kindly provided by Dr. Eric Vivier. *Il10*^fl/fl^ mice were kindly provided by Dr. Axel Roers. All experimental mice, both male and female, were 8-12 weeks old, housed in conventional housing conditions, and bred following all Institutional Animal Care and Use Committee Procedures at the Hennepin Healthcare Research Institute.

### Cytokine complex treatment

IL-15C was formed by combining 0.75 µg IL-15 (BioLegend) with 7 µg IL-15Rα-Fc (R&D Systems) and incubating for 20-30 min at 37°C prior to injection. Mice were treated on days -3 and -5 relative to infection, immunization, or harvest.

### Plasmodium infections

*Plasmodium berghei* ANKA (PbA) and PbNK65 were passaged *in vivo*, and stocks frozen in Alsever’s solution and glycerol (9:1 ratio) were stored in liquid nitrogen until use. Freshly thawed stocks of PbA or PbNK65-infected RBCs (1 x 10^6^) were diluted in PBS and injected i.v. into experimental mice.

### Anti-malarial drug treatments

For harvests after 7 dpi, mice were given water containing 200 µg/mL chloroquine (Sigma) to drink a*d libitum* from days 7-14 pi. Additionally, on days 7-9 pi, mice were treated 100 µg artesunate (Sigma) diluted in PBS and delivered i.p. in 100 µl.

### Peptide-adjuvant immunization

Mice were immunized with 100 µg ETRAMP peptide, 50 µg CD40 agonist Ab (clone FGK4.5; Bio X Cell), and 50 µg polyinosinic:polycytidylic acid [poly(I:C), Biotechne]. All vaccinations were prepared by mixing each component in PBS and were injected in 100 µl.

### Anti-IL-10R Ab treatment

Mice were treated with 200 µg of anti-IL-10R Ab (clone 1B1.3A, Bio X Cell) or a control rat IgG1 Ab (clone HRPN, Bio X Cell) on days -1, +1, and +3 relative to PbA infection. Antibodies were diluted in PBS and injected in 200 µl.

### Flow cytometry

Mice were euthanized by CO2 overdose then spleens and lymph nodes (where noted) were removed. Spleens were mashed through 70 µm cell strainers (BD Bioscience). The cells were washed and resuspended in FACS buffer (PBS with 2% FBS). ETRAMP-specific CD4^+^ T cells were enriched using APC- and PE-conjugated tetramers and EasySep APC and PE Positive Selection Kits (STEMCELL Technologies). Cells were then stained with fluorescent dye-conjugated antibodies (see **Table I**) and fixed (Foxp3 fixation/permeabilization Kit, Invitrogen). Intracellular stains (i.e., transcription factors) were performed using 1X permeabilization buffer (Foxp3 fixation/permeabilization Kit, Invitrogen). Samples were acquired on a BD LSRII Fortessa using BD FACSDiva (BD Bioscience), and data were analyzed with FlowJo v9 software (Tree Star Technologies).

### PbA ELISA

High-binding microtiter plates were coated with PbA lysate (10 µg/mL) in PBS for 1 hour incubation at 37°C. After washing with 0.5% Tween 20 (Sigma) in PBS, plates were blocked with 1% bovine serum albumin (BSA, Sigma) for 1-2 hours at room temperature. Sera samples were diluted in dilution buffer (0.2% BSA and 0.05% Tween 20 in PBS). After washing, plates were incubated with sera samples in serial dilutions for 1 hour at 37°C. The plates were washed and incubated with a peroxidase-conjugated goat anti-mouse anti-IgM or anti-IgG antibody (Southern Biotech). After washing, plates were incubated with tetramethyl-benzidine developing reagent (Surmodics) followed by addition of a stop solution. Absorbance was read at 450 nm on a Biotek Eon plate reader. Sera samples from five uninfected mice were included in each assay. The OD values from the uninfected samples were averaged, and the infected sample OD values were calculated as a fold change.

### Statistical analysis

All data were analyzed on GraphPad Prism 8. Data were evaluated for statistically significant differences using a two-tailed, unpaired *t* test with or without Welch’s correction, a one-way analysis of variance (ANOVA) test followed by Tukey’s multiple comparison test, a two-way ANOVA followed by a Bonferroni multiple comparison test, or a log-rank (Mantel-Cox) test. A *P*-value < 0.05 was considered statistically significant. All differences not specifically indicated to be significant were not significant (*P* > 0.05).

## RESULTS

### Novel peptide:MHCII tetramer detects Plasmodium-specific CD4^+^ T cells

CD4^+^ T helper (Th) cells are critical for the effective induction of humoral immunity to most pathogens and vaccines. Thus, identifying antigen-specific CD4^+^ T cells is key to define the immune response following natural infection and determine how this response is affected by therapies. The lack of parasite-derived T cell epitopes has impaired the study of antigen-specific CD4^+^ T cell responses against *Plasmodium*. Instead, groups have utilized surrogate activation markers (14), transgenic malaria parasites expressing model antigens (21, 22), or TCR transgenic mice specific for *Plasmodium* antigens (23, 24). These strategies have resulted in important findings regarding the development and function of antigen-specific CD4^+^ T cell responses during *Plasmodium* infection in mice. However, these systems do not permit the analysis of endogenous antigen-specific CD4^+^ T cells responding to *Plasmodium* at a natural precursor frequency. Draheim and colleagues recently identified MHC class II-binding peptides in *Plasmodium berghei* ANKA (PbA) (25). We tested the immunodominant peptide sequences in an I-A^b^ binding algorithm and identified a strong I-A^b^-binding peptide from the early transcribed membrane protein (ETRAMP276-286) in PbA: NYSIPRPNVTS. We generated a peptide:MHCII tetramer that successfully identifies peptide-specific CD4^+^ T cells in C57BL/6 mice following PbA infection (**Fig. 1**).

**FIGURE 1.**
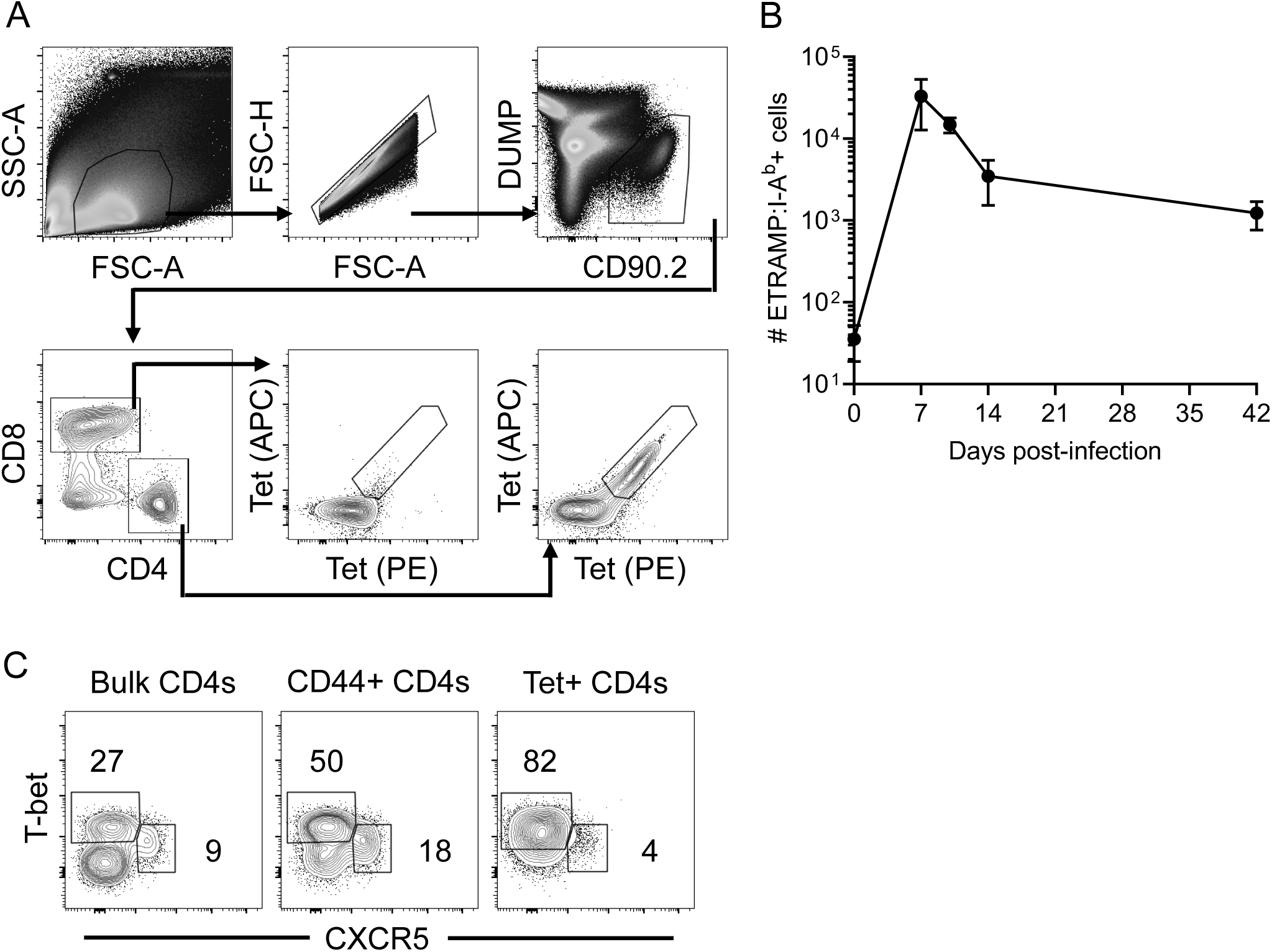
ETRAMP:I-A^b^ tetramer detects PbA-specific CD4^+^ T cells. At 7 dpi, spleen cells were stained with ETRAMP:I-A^b^ tetramer conjugated to APC and PE and enriched using magnetic bead-based technology. (A) Gating strategy: after gating on lymphocytes, single cells, and live CD19^-^ F4/80^-^ CD90.2^+^ cells, ETRAMP:I-A^b^ tetramer-binding CD4^+^ T cells were identified. Few ETRAMP:I-A^b^ tetramer-binding cells were found in the CD8^+^ T cell gate. (B) Number of ETRAMP:I-A^b^ tetramer-binding cells at 0, 7, 10, 14, and 42 dpi. Data are compiled from 6-14 mice per timepoint and displayed as mean ± SD. (C) Representative flow plots demonstrating T-bet and CXCR5 expression on bulk CD4^+^ T cells, CD44^+^ CD4^+^ T cells, and tetramer-binding CD4^+^ T cells.

Using an established magnetic enrichment protocol method (26), we quantified the number of ETRAMP-specific CD4^+^ T cells over the course of PbA infection in mice to track expansion and contraction kinetics as well as define the phenotype of these cells. After gating on lymphocytes, we excluded doublets as well as CD19^+^ B cells, F4/80^+^ macrophages and dead cells (“Dump” gate). Next, we gated on CD90.2^+^ T cells, then CD4^+^ CD8^-^ T cells. Using the tetramer in APC and PE, we identified dual tetramer-binding cells (**Fig. 1A**). Uninfected adult mice (8-12 weeks old) had on average 35 ETRAMP-specific CD4^+^ T cells in lymphoid tissues (spleen and lymph nodes), which is similar to the precursor frequency of other highly-studied CD4^+^ T cell epitopes (26). We found that ETRAMP-specific CD4^+^ T cell expansion peaks at about 7 days post-infection (dpi), when the number of ETRAMP-specific CD4^+^ T cells is approximately 1,000-fold higher than the precursor frequency (**Fig. 1B**). The ETRAMP-specific CD4^+^ T cell response quickly contracts, likely due to antigen clearance since mice are treated with anti-malarial drugs starting at 7 dpi (**Fig. 1B**). At 42 dpi, the number of ETRAMP-specific CD4^+^ T cells is approximately 50-fold higher than the precursor frequency, indicating a stable memory population (**Fig. 1B**). Similar to other groups (12, 28, 29), we found that PbA infection is dominated by a Th1 response, as determined by T-bet staining, with very few CXCR5^+^ Tfh cells among the ETRAMP-specific CD4^+^ T cells at 7 dpi (**Fig. 1C**). Overall, these data demonstrate that this ETRAMP:I-A^b^ tetramer is a novel tool to identify *Plasmodium*-specific CD4^+^ T cells and test strategies to modulate that response.

### IL-15C treatment increases the frequency and number of Plasmodium-specific Tfh cells at 7 days post-infection

We previously showed that IL-15C treatment induces IL-10 production from NK cells, which protects against PbA-induced experimental cerebral malaria (27). In addition to TCR and co-stimulation, cytokines play critical roles in directing CD4^+^ T cell differentiation and function (30), suggesting that dysregulated inflammation could impair appropriate CD4^+^ T cell differentiation. IL-10 may have particular utility due to its ability to dampen the pathologic Th1-driven immune response during PbA infection. Therefore, we investigated the effect of IL-15C treatment on the phenotype of ETRAMP-specific CD4^+^ T cells. We found that IL-15C treatment prior to infection had no effect on the total number of ETRAMP-specific CD4^+^ T cells in the spleen at 7 dpi (**Fig. 2A**). However, IL-15C treatment resulted in increased frequency and numbers of ETRAMP-specific CXCR5^+^PD-1^+^ Tfh cells in the spleen at 7 dpi (**Fig. 2B-D**) as well as polyclonal (bulk) CD4^+^ T cells (**Fig. 2E-F**). All ETRAMP-specific CXCR5^+^PD-1^+^ cells were Bcl-6^+^ and T-bet^-^. IL-15C treatment resulted in a slight but significant decrease in the frequency and number of ETRAMP-specific T-bet^+^ Th1 cells in the spleen at 7 dpi (**Supplemental Fig. 1A-B**) but had no effect on the frequency or number of ETRAMP-specific Foxp3^+^ Treg cells at 7 dpi (**Supplemental Fig. 1C-D**).

**FIGURE 2.**
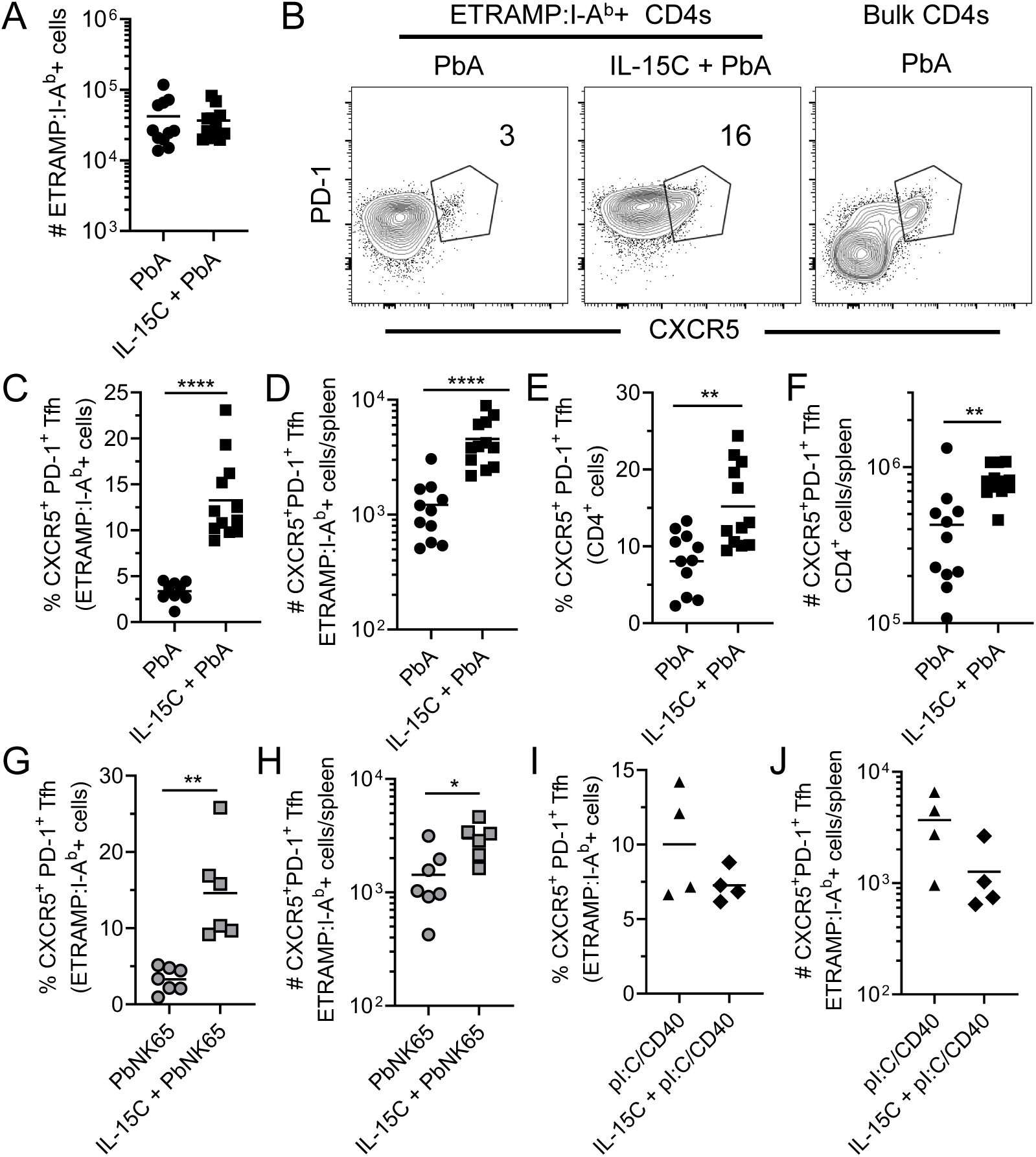
IL-15C treatment results in increased ETRAMP:I-A^b^-specific CXCR5^+^PD-1^+^ CD4^+^ Tfh cells at 7 days following PbA and PbNK65 infection. Mice were left untreated or treated with IL-15C prior to PbA infection (A-F), PbNK65 infection (G-H), or immunization with ETRAMP peptide, anti-CD40 agonist and polyI:C (I-J). ETRAMP-specific CD4^+^ T cells were enriched from spleens of mice at 7 dpi and assessed by flow cytometry. (A) Total number of ETRAMP-specific CD4^+^ T cells. (B) Representative flow plots of CXCR5 and PD-1 staining on ETRAMP-specific CD4^+^ T cells or bulk (polyclonal) CD4^+^ T cells. Frequency (C, G, I) and number (D, H, J) of ETRAMP-specific CXCR5^+^PD-1^+^ Tfh cells. Frequency (E) and number (F) of polyclonal CXCR5^+^PD-1^+^ Tfh cells. Symbols represent individual mice; horizontal bars indicate mean values. Data compiled from three (A-F) or two (G-H) independent experiments or from one experiment (I-J). * *P* < 0.05, ** *P* < 0.01, **** *P* < 0.0001 as determined by two-sided *t* test or Mann-Whitney U test.

PbNK65 is genetically similar to PbA but does not cause a cerebral malaria-like disease in C57BL/6 mice (31, 32). The ETRAMP peptide sequence is also encoded in PbNK65, allowing use of our tetramer to investigate the antigen-specific CD4^+^ T cell response to PbNK65. We found that IL-15C treatment prior to PbNK65 infection also resulted in increased frequency and number of ETRAMP-specific CXCR5^+^PD-1^+^ Tfh cells at 7 dpi (**Fig. 2G-H**). Thus, IL-15C modulates CD4^+^ T cell differentiation, promoting a Tfh fate, in various *Plasmodium* infection models.

A vaccination strategy that has been effective at inducing strong T cell immunity and a Th1-dominated response is immunizing with an anti-CD40 agonist antibody along with polyI:C and a peptide of interest (33). Antigen-specific CD8^+^ T cell and CD4^+^ T cell responses to this immunization strategy have been assessed (33, 34). We tested whether IL-15C treatment induced stronger Tfh differentiation in the context of this immunization strategy. We found that IL-15C treatment prior to CD40/polyI:C/peptide immunization had no impact on the frequency or number of ETRAMP-specific CXCR5^+^PD-1^+^ Tfh cells (**Fig. 2I-J**). Overall, these data suggest that *Plasmodium* infection creates a unique inflammatory environment that can be modulated by IL-15C treatment to promote Tfh differentiation.

### Genetic deletion of NK cell-derived IL-10 prevents IL-15C-induced Tfh differentiation

We previously showed that IL-15C treatment induces IL-10 production from NK cells and that IL-10 expression by NK cells was required for IL-15C-mediated survival following PbA infection (27). We next sought to investigate whether IL-10 expression by NK cells was necessary for enhanced Tfh differentiation following IL-15C treatment. To test this requirement, we bred IL-10^fl/fl^ mice to NKp46-iCre mice (35) to genetically delete IL-10 from NK cells (referred to as NK*^Il10^*^-KO^ mice). We previously showed that both IL-10 production and NK cell expansion in response to IL-15C treatment are dependent on NK cell intrinsic STAT3 signaling (36). To test if NK cell-specific loss of IL-10 expression had an impact on NK cell and/or CD8^+^ T cell expansion following IL-15C treatment, we quantified the number of splenic NK cells and CD8^+^ T cells in IL-15C-treated WT and NK*^Il10^*^-KO^ mice and found similar expansion (**Supplemental Fig. 2**). To investigate the impact of NK cell-derived IL-10 on Tfh formation, WT and NK*^Il10^*^-KO^ mice were left untreated or treated with IL-15C prior to PbA infection. At 7 dpi, the number of ETRAMP-specific CD4^+^ T cells was similar between IL-15C-treated WT and NK*^Il10^*^-KO^ mice (**Fig. 3A**), indicating that a lack of NK cell-derived IL-10 had no impact on PbA-specific CD4^+^ T cell expansion at an acute timepoint. However, while IL-15C-treated WT mice had increased Tfh formation, IL-15C-treated NK*^Il10^*^-KO^ mice had similar frequency and number of ETRAMP-specific CXCR5^+^PD-1^+^ Tfh cells as untreated WT mice (**Fig. 3B-C**). These data indicate that NK cell-derived IL-10 is required for IL-15C-mediated Tfh differentiation.

**FIGURE 3.**
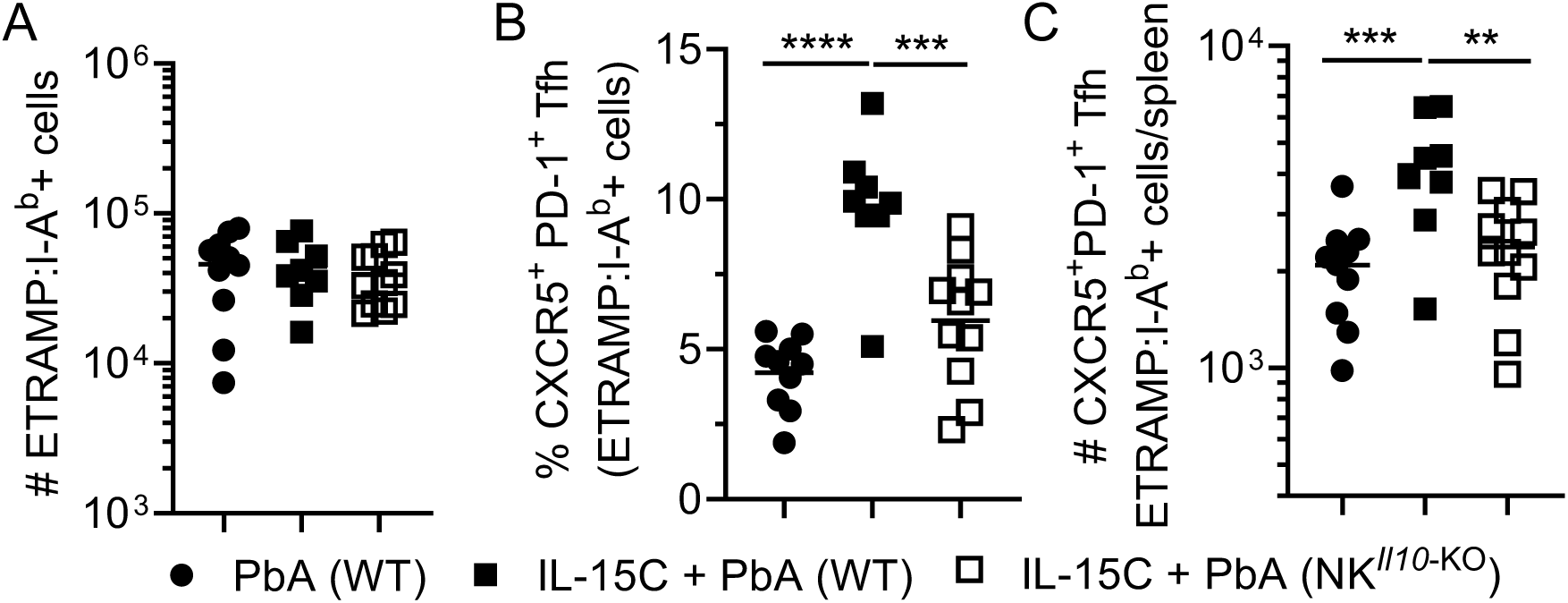
Genetic deletion of NK cell-derived IL-10 prevents IL-15C-mediated Tfh differentiation. WT (B6, IL-10^fl/fl^ or NKp46-Cre) mice or NK*^Il10^*^-KO^ mice were left untreated or treated with IL-15C prior to PbA infection. Spleens were harvested at 7 dpi. (A) Total number of ETRAMP-specific CD4^+^ T cells. Frequency (B) and number (C) of CXCR5^+^PD-1^+^ ETRAMP-specific Tfh cells. Symbols represent individual mice; horizontal bars indicate mean values. Data compiled from four independent experiments. ** *P* < 0.01, *** *P* < 0.001, as determined by one-way ANOVA followed by Tukey’s multiple comparisons test.

### Antibody blockade or genetic deletion of IL-10R on T cells prevents IL-15C-induced Tfh differentiation

We next sought to determine whether IL-10R signaling was required for IL-15C-mediated Tfh differentiation. First, mice were treated with IL-15C along with an anti-IL-10R blocking antibody or an isotype control antibody, and the phenotype of ETRAMP-specific CD4^+^ T cells was assessed at 7 dpi. Blocking IL-10R slightly increased the number of ETRAMP-specific CD4^+^ T cells (**Fig. 4A**) but reduced the frequency of ETRAMP-specific CXCR5^+^PD-1^+^ Tfh cells relative to mice treated with IL-15C and an isotype control antibody (**Fig. 4B**). Since blocking IL-10R increased the number of total ETRAMP-specific CD4^+^ T cells (**Fig. 4A**), the number of ETRAMP-specific CXCR5^+^PD-1^+^ Tfh cells was not reduced in that group (**Fig. 4C**).

**FIGURE 4.**
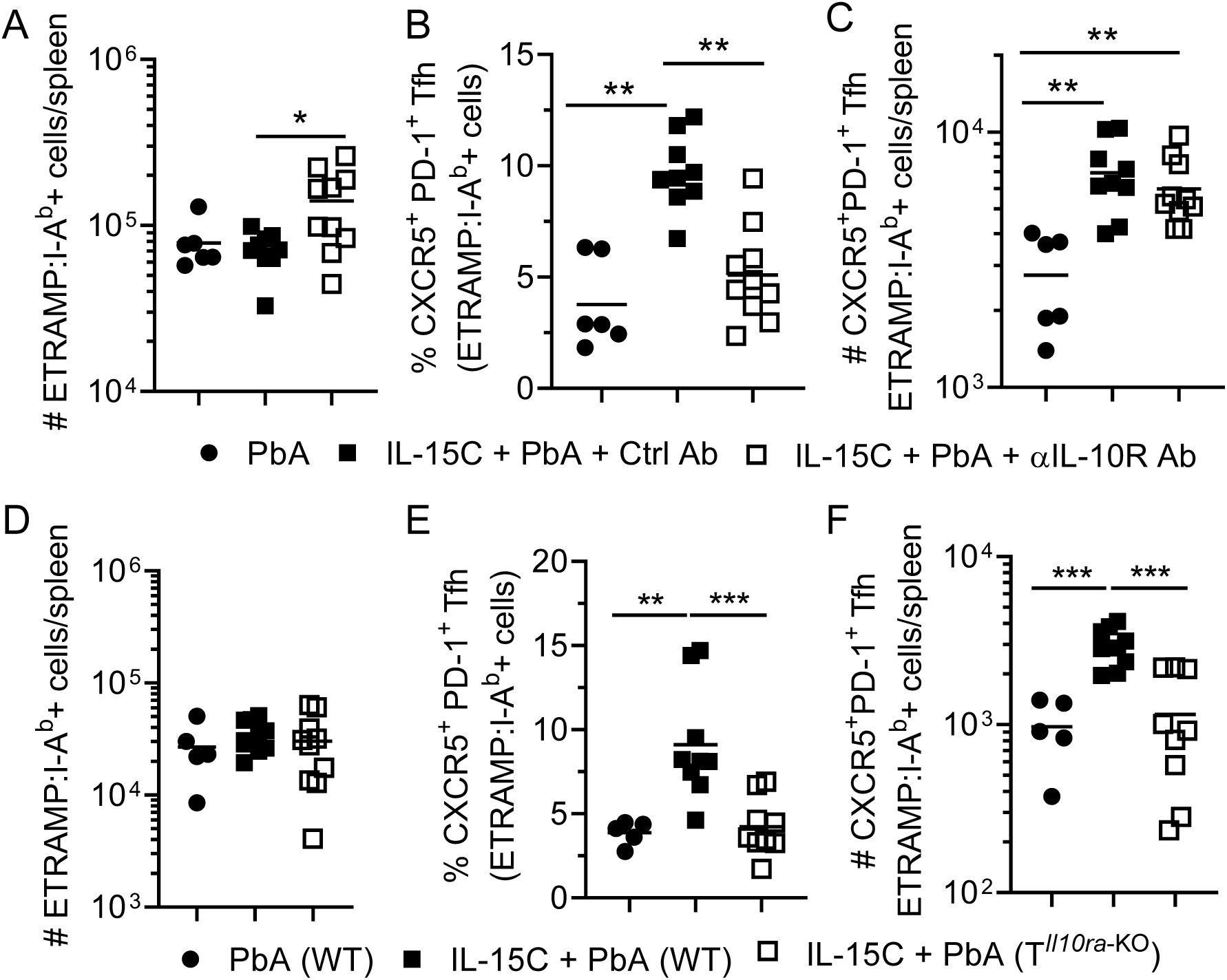
IL-10R blockade or genetic deletion of IL-10R from T cells prevents IL-15C-mediated Tfh differentiation. (A-C) WT mice were left untreated or treated with IL-15C on days -3 and -5 relative to PbA infection. On days -1, +1 and +4, mice were given 200 µg of an anti-IL-10Rα Ab or an isotype control Ab. Spleens were harvested at 7 dpi. (A) Total number of ETRAMP-specific CD4^+^ T cells. Frequency (B) and number (C) of CXCR5^+^PD-1^+^ ETRAMP-specific Tfh cells. (D-F) WT mice or T*^Il10ra^*^-KO^ mice were left untreated or treated with IL-15C prior to PbA infection. Spleens were harvested at 7 dpi. (D) Total number of ETRAMP-specific CD4^+^ T cells. Frequency (E) and number (F) of CXCR5^+^PD-1^+^ ETRAMP-specific Tfh cells. Symbols represent individual mice; horizontal bars indicate mean values. Data compiled from three (A-C) or two (D-F) independent experiments. * *P* < 0.05, ** *P* < 0.01, as determined by one-way ANOVA followed by Tukey’s multiple comparisons test.

Multiple cell types express the IL-10 receptor, complicating the interpretation of systemic IL-10R blockade. To determine whether IL-10R signaling in CD4^+^ T cells was regulating Tfh differentiation, we crossed CD4-Cre mice to IL-10Rα^fl/fl^ mice to generate mice that lack IL-10Rα expression on T cells (CD4*^Il10ra^*^-KO^ mice). The number of ETRAMP-specific CD4^+^ T cells was similar between IL-15C-treated WT and CD4*^Il10ra^*^-KO^ mice (**Fig. 4D**). However, while IL-15C treatment of WT mice increased Tfh formation, IL-15C-treated CD4*^Il10ra^*^-KO^ mice showed no increase in the frequency or number of ETRAMP-specific CXCR5^+^PD-1^+^ Tfh cells (**Fig. 4E-F**). These data indicate that IL-10R expression on T cells is required for IL-15C-enhanced Tfh differentiation. Collectively, we show that IL-15C-induced expression of IL-10 from NK cells acts directly on T cells to promote Tfh differentiation in the context of *Plasmodium* infection.

### IL-15C treatment results in increased Plasmodium-specific IgG at 42 days post-PbA infection and improves survival following reinfection

Since dysregulated inflammation during *Plasmodium* infection disrupts the formation of GCs, resulting in decreased antibody production (12), and IL-15C treatment promotes Tfh differentiation, we hypothesized that IL-15C may result in increased antibody production at a memory time point. To test this premise, mice were left untreated or treated with IL-15C prior to PbA infection. At 7 dpi, all mice were treated with the anti-malarial drugs artesunate and chloroquine to clear the infection. At either 10 or 42 dpi, anti-PbA IgM and IgG antibody levels in the serum were quantified via ELISA. At 10 dpi, untreated and IL-15C-treated mice had similar PbA-specific IgM and IgG levels (**Fig. 5A-B**). However, at 42 dpi, IL-15C-treated mice had similar PbA-specific IgM levels as untreated mice but significantly increased PbA-specific IgG levels (**Fig. 5C-D**). In addition, we found that IL-15C-treated NK*^Il10^*^-KO^ had significantly reduced PbA-specific IgG compared with IL-15C-treated WT mice at 42 dpi (**Fig. 5D**). Together, these findings demonstrate that IL-15C treatment, operating through NK cell-derived IL-10, induces Tfh differentiation and enhances antibody production.

**FIGURE 5.**
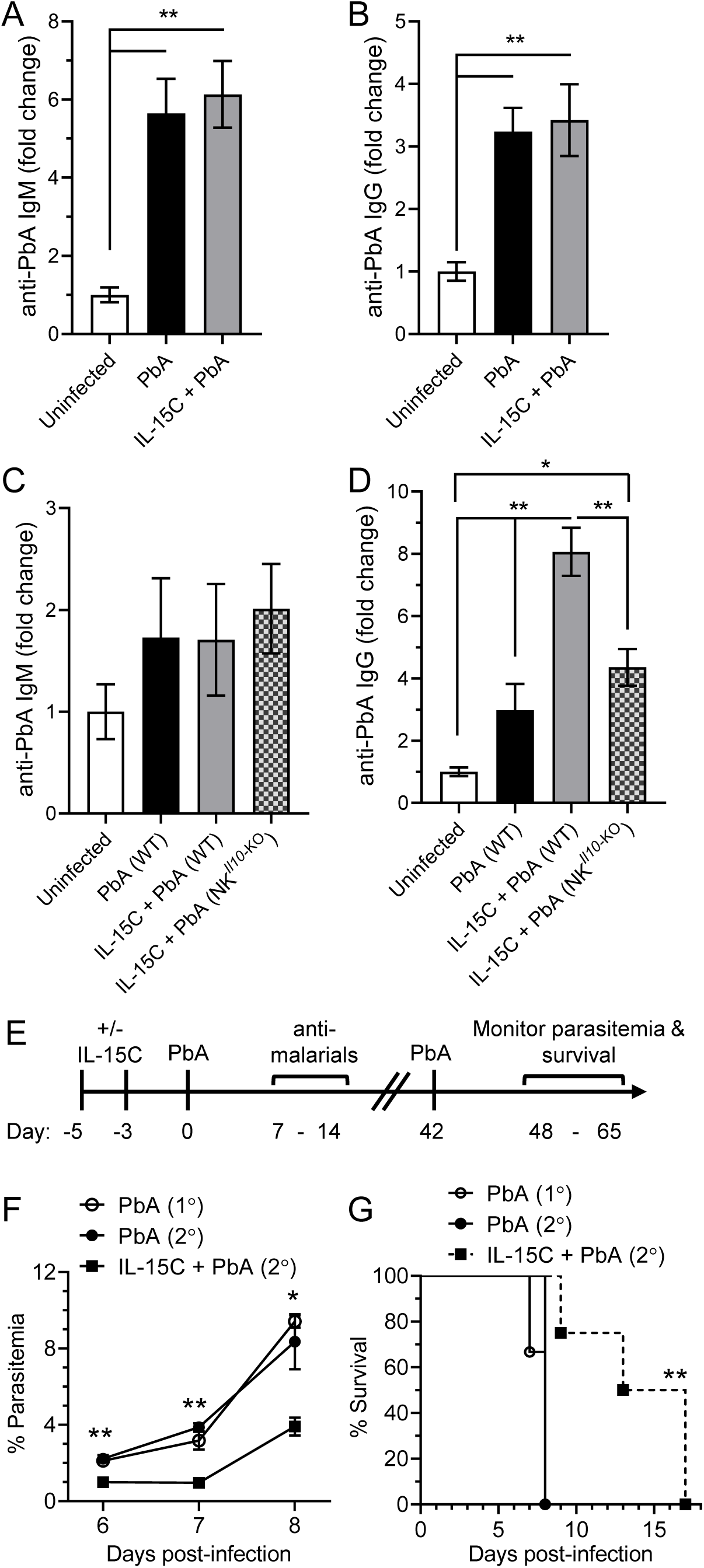
IL-15C-treated mice have increased PbA-specific IgG at 42 dpi and are better protected from reinfection. (A-D) Mice were either left untreated or treated with IL-15C prior to PbA infection. At 10 dpi (A-B) or 42 dpi (C-D), PbA-specific IgM (A and C) or IgG (B and D) were quantified via ELISA. * *P* < 0.05, ** *P* < 0.01, as determined by one-way ANOVA followed by Tukey’s multiple comparisons test. Data compiled from two independent experiments per timepoint with a total of 7-12 mice per group and displayed as mean ± SEM. (E-G) Mice were either left untreated (n = 6) or treated with IL-15C (n = 4) prior to PbA infection; at 7 dpi, mice were treated with anti-malarial drugs. At 6 weeks post-infection, mice were reinfected with PbA (2°). A group of mice (n = 3) were infected with PbA for the first time (1°) as a control for the development of experimental cerebral malaria. (E) Schematic of experimental design. (F) Parasitemia was determined via blood smear from days 6-8 pi. ** *P* < 0.01, as determined by one-way ANOVA followed by Tukey’s multiple comparisons test on data from 6 and 7 dpi. * *P* < 0.05, as determined on two-tailed *t* test on data from 8 dpi, comparing the PbA (2°) and IL-15C + PbA (2°) groups. (G) Survival curve. ** *P* < 0.01, as determined by Log-rank (Mantel-Cox) test between the PbA (2°) and IL-15C + PbA (2°) groups.

Others have demonstrated that mice can be reinfected with PbA multiple times, with three reinfections needed before improved survival is observed (37). To test if IL-15C treatment prior to a primary (1°) PbA infection affects the outcome of a secondary (2°) PbA infection, mice were either left untreated or treated with IL-15C prior to PbA infection (**Fig. 5E**). At 7 dpi, mice were treated with anti-malarial drugs, and at 6 weeks post-infection, mice were reinfected with PbA (2° infection). A group of mice were infected with PbA for the first time (1° infection) as a control for the development of experimental cerebral malaria. Mice treated with IL-15C prior to their first PbA infection had significantly lower parasitemia levels on days 6, 7, and 8 pi following reinfection (**Fig. 5F**). Additionally, IL-15C treatment resulted in a significant improvement in survival following reinfection (**Fig. 5G**). In sum, these data indicate that IL-15C treatment prior to PbA infection results in improved immunity to a secondary infection.

## DISCUSSION

NK cells have the dual ability to produce pro-inflammatory (IFN-γ) and anti-inflammatory (IL-10) cytokines, demonstrating their crucial role both as inflammatory effectors for infection clearance and as regulators to limit immune-mediated damage to the host. This balance between elimination of infectious pathogens while limiting pathology is particularly relevant in systemic infections, such as *Plasmodium*, when pathogens that have not been contained can lead to a dysregulated systemic inflammatory response, leading to morbidity or mortality (38). Indeed, high IL-10 levels are detected in the plasma of children with severe malaria (39–41), suggesting that IL-10 is induced in an attempt to dampen the pathologic systemic inflammation. IL-10-producing NK cells have been described in the context of various infections (42–45), including PbA (27). We previously identified a method to specifically induce IL-10 expression in mouse and human NK cells using IL-15 (27). Moreover, we demonstrated that these immunoregulatory IL-10-producing NK cells dampen the pathologic inflammatory response, including IFN-γ production, during PbA infection (27). Interestingly, IL-10 production correlates with ongoing but asymptomatic *Plasmodium* infection in young children (46), suggesting that IL-10 may mitigate disease symptoms during ongoing parasite exposure. Additionally, studies in humans and infection models revealed that inflammatory cytokines that contribute to the induction of symptomatic malaria, such as IFN-γ, also upregulate the expression of T-bet in Tfh cells, which impairs their capacity to provide help for antibody responses (11, 12). These data suggest a model whereby IL-10-producing NK cells can be induced at the appropriate time and anatomic location to both limit disease symptoms and modulate the inflammatory milieu to promote Tfh differentiation and antibody production.

Antibodies are a key component of natural and vaccine-derived immunity to *Plasmodium.* CD4^+^ T helper cells, especially Tfh cells, which co-localize with B cells in GCs, promote long-lived and high-quality antibody responses. However, malaria immunity is relatively short in duration and rapidly wanes in the absence of re-infection. The mechanisms underlying the slow acquisition and subsequent decay of anti-*Plasmodium* immunity remain incompletely understood, but dysregulated inflammation is thought to impair immunity (11–16). *Plasmodium*-specific CD4^+^ T cells are a correlate of protective immunity following either natural exposure or vaccination (47–50). However, several studies have identified pathologic roles for Th1 cells and IFN-γ during *Plasmodium* infection (11–14). Indeed, the dysregulated inflammatory response to *Plasmodium* infection disrupts the GC reaction. In *Plasmodium*-infected children, a Th1-polarized response negatively affects the differentiation of GC B cells (11). *P. falciparum* infection of *Saimiri sciureus* monkeys results in the disruption of GC architecture (15). In mice, Th1 cytokines (IFN-γ, TNF-α) suppress GC responses during *Plasmodium* infection (12, 13), and in the absence of properly activated Tfh cells, there is a complete disruption of the GC response and an inability to clear *Plasmodium* (16). Thus, a better understanding of the *Plasmodium*-specific CD4^+^ T cell response would inform strategies to boost Tfh formation and resulting antibody production.

We generated a new pMHCII ETRAMP tetramer to identify and analyze antigen-specific CD4^+^ T cells responding to *Plasmodium* infection at a natural precursory frequency. Similar to others (12, 28, 29), we found that PbA infection is dominated by a Th1 response, with very few Tfh cells at 7 dpi (Fig. 1). Interestingly, IL-15C treatment boosted ETRAMP-specific Tfh formation in the context of both PbA and PbNK65 infection but not in the context of a peptide-adjuvant vaccination model (Fig. 2). These data suggest that the immune response to *Plasmodium* infection may be uniquely susceptible to manipulation by IL-15C treatment. Furthermore, using NK cell specific genetic knockout mice, we demonstrated that IL-10 production from NK cells was required for IL-15C-mediated Tfh differentiation (Fig. 3). To further confirm that IL-10 signaling was required for Tfh differentiation, mice were treated with IL-15C along with an anti-IL-10R blocking antibody. We found that blocking IL-10R resulted in a reduced frequency of ETRAMP-specific Tfh cells at 7 dpi (Fig. 5). These data suggest that blocking IL-10 signaling prevents IL-15C-induced Tfh differentiation but did not identify the cell type(s) responding to NK cell-derived IL-10.

IL-10 has pleiotropic effects including reducing IFN-γ and/or IL-12 production (51–55), which may indirectly affect Tfh differentiation. Moreover, many cell types express the IL-10 receptor, including CD4^+^ T cells. Notably, IL-10 signals through STAT3, which is required for CXCR5 expression and Tfh differentiation (56). Thus, we reasoned that CD4^+^ T cells may respond directly to NK cell-derived IL-10. Indeed, IL-15C failed to increase Tfh differentiation in mice lacking IL-10R expression selectively on T cells (Fig. 4). While these data don’t rule out a role for NK cell-derived IL-10 modulating other immune cells during PbA infection, these data demonstrate that direct IL-10 signaling in T cells is important for Tfh differentiation following IL-15C treatment. Our data provide insight into the interplay between IL-10^+^ NK cells and T cells, which is particularly relevant as strategies to target and use NK cells for adoptive therapies are increasingly developed for cancer therapies and vaccines (57–59).

PbA infection results in a Th1-dominated immune response that dampens GC formation, GC B cell differentiation, and antibody production (12, 60). We found that IL-15C-treated mice had higher PbA-specific IgG levels, which was dependent on NK cell-derived IL-10 (Fig. 5). During *P. yoelii* infection, B cell-intrinsic IL-10 signaling is required for GC B cell reactions, isotype-switched antibody responses, parasite control, and host survival (13), and CD4^+^ T cells are a critical source of IL-10 in *P. yoelii* infection (61). Future studies will investigate the impact of IL-10 signaling on B cell differentiation, GC formation, and antibody production in the context of PbA infection. We also showed that mice treated with IL-15C prior to a primary PbA infection had reduced parasitemia and improved survival following a secondary PbA infection (Fig. 5). While antibodies are important for controlling *Plasmodium* infection (62), CD4^+^ T cells also play critical roles in controlling parasitemia in mouse models of *Plasmodium* infection (24, 63–66). Thus, future studies will investigate the effect of IL-15C treatment on antibody affinity maturation and effector functions as well as memory CD4^+^ T cell recall responses and their roles in improving survival following reinfection.

Overall, our data demonstrate that IL-15C – via its induction of IL-10 from NK cells – modulates the dysregulated immune response during *Plasmodium* infection to promote Tfh differentiation and increase antibody production. Further studies are needed to better understand the impact of NK cell-derived IL-10 on CD4^+^ T cell differentiation and memory formation and downstream effects on GC formation, B cell differentiation, and antibody production. In sum, these data provide important insight into the role of dysregulated inflammation during blood-stage *Plasmodium* infection and identify pathways that could be exploited to enhance immunity to *Plasmodium* as well as other infections or inflammatory diseases.

## ACKNOWLEDGEMENTS

We would like to thank Dr. Marc Jenkins for his scientific input on this manuscript. We would like to thank Dr. Adam Burrack for assistance with the I-A^b^ peptide algorithm to identify the ETRAMP peptide.

## Notes

### Competing Interest Statement

The authors have declared no competing interest.

