## Supplemental Figures for "IL-15 complex-induced IL-10 enhances *Plasmodium*-specific CD4^+^ Tfh differentiation and antibody production"

SUPPLEMENTAL FIGURE 1

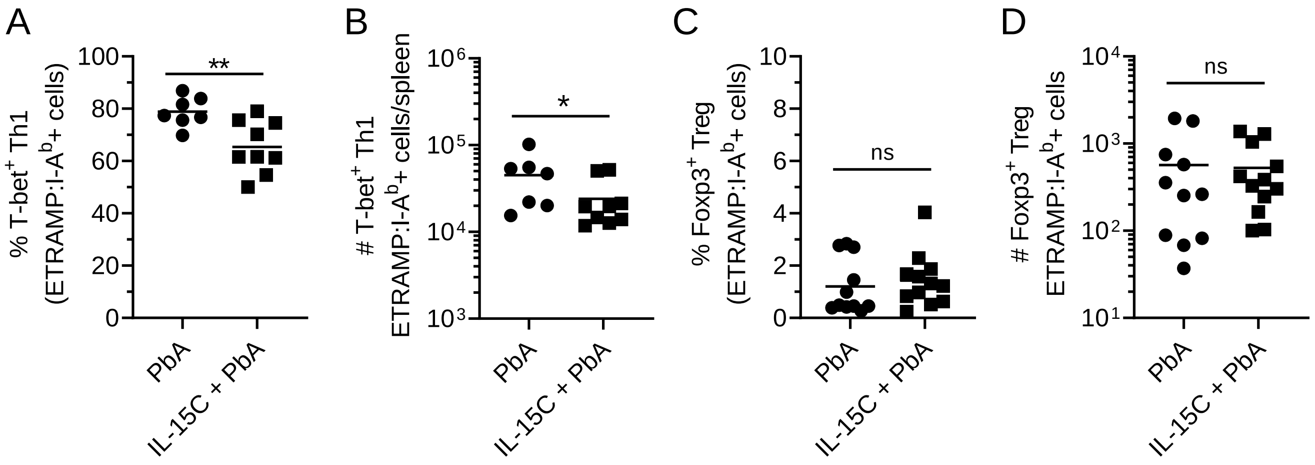


**SUPPLEMENTAL FIGURE 1. IL-15C treatment results in decreased ETRAMP:I-A$^{b}$-specific T-bet$^{+}$ CD4 Th1 cells at 7 dpi.** Mice were left untreated or treated with IL-15C prior to PbA infection. ETRAMP-specific CD4 T cells were enriched from spleens of mice at 7 dpi and assessed by flow cytometry. Frequency (A, C) and number (B, D) of ETRAMP-specific T-bet$^{+}$ Th1 cells (A, B) and Foxp3$^{+}$ Treg cells (C, D). Symbols represent individual mice; horizontal bars indicate mean values. Data compiled from 3 independent experiments. * $P < 0.05$, as determined by Mann-Whitney U test. ns = not significant.

### SUPPLEMENTAL FIGURE 2

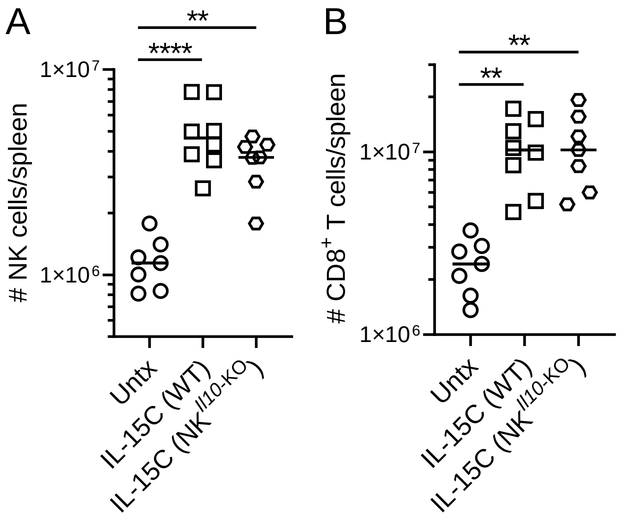


**SUPPLEMENTAL FIGURE 2. IL-15C-induced expansion of NK cells and CD8$^{+}$ T cells is not impacted by genetic deletion of NK cell-derived IL-10.** WT or NK$^{Il10\text{-KO}}$ mice were left untreated or treated with IL-15C. Spleens were harvested three days after the second IL-15C treatment and analyzed by flow cytometry. (A) Number of live, CD3$^{-}$ NK1.1$^{+}$ NKp46$^{+}$ NK cells. (B) Number of live, CD3$^{+}$ CD8$^{+}$ T cells. Symbols represent individual mice; horizontal bars indicate mean values. Data compiled from two independent experiments. ** $P < 0.01$, **** $P < 0.0001$, as determined by one-way ANOVA followed by Tukey’s multiple comparisons test.
